# Interindividual variability in HRV reactivity: autonomic flexibility or regression to the mean?

**DOI:** 10.1101/274118

**Authors:** Dimitriy A. Dimitriev, Elena V. Saperova, Olga S. Indeykina, Aleksey D. Dimitriev

## Abstract

**Objective:** There is now a substantial body of evidence linking the baseline level of heart rate variability (HRV) with the magnitude of stress-induced reduction in respiratory sinus arrhythmia. However, it remains to be proved to what extent these interindividual differences in stress responses may be attributed to the statistical phenomenon of regression to the mean (RTM). We sought to test the hypothesis that the statistical artifact RTM explains part of the baseline effect.

**Approach:** Heart rate recording was carried out in 1,156 volunteers. To obtain an estimate of the stress response, 148 persons were randomly selected. Participants were monitored on a rest day and just before an academic examination for state anxiety and HRV. Participants were divided into quartiles according to baseline HRV levels and were compared for response to academic stress.

**Main results:** We observed a significant reduction in HRV in subjects with a high baseline HRV (> 75th percentile), while a significant increase was found in the group with low baseline HRV. Regression analysis demonstrated that the value of baseline HRV correlated with the magnitude of stress reaction consistent with the RTM model. Baseline-adjusted ANCOVA does not reveal significant intergroup differences in the changes in heart rate (HR) and HRV from rest to exam. RTM-adjusted estimates confirmed an exam effect for HR and HRV.

**Significance:** The results of our study strongly support RTM as the source of variability of stress-related changes in HRV.

## 1. Introduction

Although the precise nature of emotion is not yet clear, theorists and researchers agree that emotion involves changes across multiple response systems (Gross et al. 2006). According to Cohen et al. (1995), psychological stress occurs when an individual perceives that the situation taxes or exceeds his or her adaptive capacity to cope with it. The model of neurovisceral integration has been widely accepted as a heuristically useful framework for the study of psychological stress (Thayer and Lane 2000). This model conceptualizes that autonomic flexibility provides the basis for adaptation to environmental challenge. Autonomic flexibility can be described as the capacity of the autonomic nervous system (ANS) to adapt to changes in the environment by modifying arousal, respiration, heart rate (HR), and attention (Friedman and Thayer 1998). Porges’ proposed polyvagal theory (Porges 1995) postulates that smart vagus (nucleus ambiguous branch of the parasympathetic nervous system) provides the neurophysiological substrate for social communication, attention, and self-regulation (Sinnreich et al. 1998).

Respiratory sinus arrhythmia (RSA), the rhythmical fluctuation in RR intervals accompanying breathing, has been widely used as an index of cardiac vagal tone (Berntson et al. 1993). Low rest RSA and redundant RSA withdrawal in response to emotional challenge are associated with psychopathology, anxiety, and panic disorder (Beauchaine 2001; Licht et al. 2009; Chalmers et al. 2014). These data suggest that RSA can function as a reliable indicator of emotional reactivity. Diminished autonomic flexibility refers to decrease of variability in HR and blunted RSA (Friedman and Thayer 1998). The direct manifestation of flexibility is reactivity of HR and RSA; HR and RSA reactivity can be determined as the difference between stress and rest scores. Thus, a high level of rest RSA should be associated with high RSA reactivity and baseline RSA is a predictor of emotional performance. Observations of stress-induced changes in anxiety and RSA, however, are equivocal.

Kok et al. (2010) found that vagal tone and psychosocial well-being reciprocally and prospectively predict one another. High baseline RSA was related to low trait anxiety (Miu et al. 2009) and high RSA reactivity (Rottenberg et al. 2007). In contrast, Alkozei et al. (2015) reported the absence of differences at rest or in response to stress between anxious and non-anxious children. Gerra et al. (2000) found that adolescent boys with an anxiety disorder showed a higher HR reactivity to a mixed-model stress task than controls. Thus, a higher level of autonomic flexibility was typical for anxious boys. Moreover, there are a growing number of observations that do not fit the theories of Thayer and Porges (Kossowsky et al. 2012; Gorka et al. 2013; Fortunato et al. 2013; Wang et al. 2013; Kristensen et al. 2014; Spangler et al. 2015; Couyoumdjian et al. 2016; Fung et al. 2017). One possible explanation for this disagreement could be the influence of individual differences in autonomic control and autonomic flexibility on RSA reactivity (Berntson and Cacioppo 2004).

The results of the studies outlined above have indicated that autonomic flexibility (i.e., reactivity) depends on the baseline level of RSA (Friedman and Thayer 1998; Beauchaine 2001; Rottenberg et al. 2007; Couyoumdjian et al. 2016). High baseline RSA was associated with a decline in RSA and subjects with low baseline RSA had the opposite reaction to stress. A plausible explanation of this phenomenon is regression to the mean (RTM). There is a strong theoretical rationale for this proposal. The RTM prediction is that extreme baseline levels tend to become less extreme after repeated measurements, even in the absence of real change. Previous studies have demonstrated that RTM is the consequence of random fluctuation or nonsystematic error in repeated measurement (Barnett et al. 2005). Davis (1976) has characterized RTM as the phenomenon by which “a variable that is extreme on its first measurement will tend to be close to the center of the distribution for a later measurement.”

The theoretical aspects of RTM are outlined by Lin and Hughes (1997). RTM can be identified by considering the regression (Barnett et al. 2005)

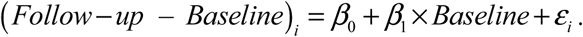

A negative regression coefficient *β*_1_ implies that subjects with very low baseline levels tend to increase and subjects with unusually high baseline levels tend to decrease. This relationship illustrates the RTM effect. Several methods have been proposed for adjusting for the effect of RTM when the variable of interest is supposed to be normally distributed (Lin and Hughes 1997).

Notwithstanding the importance of this issue, the empirical evidence on the effects of RTM on physiological measures is rather sparse and, in particular, the evidence concerning the effects of RTM on autonomic reactivity to psychosocial stress. Some authors consider RTM as a source of bias; they discuss results regarding RTM. Unfortunately, these authors did not test the hypothesis that RTM explains, at least in part, observed changes. One exception is the research of Gotfredsen et al. (1997), who tested for the RTM effect by using a modification of the method of Mee and Chua.

This study aims to evaluate the contribution of RTM and to examine implications for interpreting findings from follow-up HRV research.

## 2. Methods

### 2.1 Participants

Data for this study were obtained from research on a cohort conducted from 2005 to 2015. A total of 1,156 students (286 men/870 women, age range from 19 to 24 years (mean age (mean ± SE): 20.53 ± 0.11)) of Chuvash State Pedagogical University participated in the study of heart rate variability (HRV). Each volunteer underwent a comprehensive screening procedure that included physical examination, health history questionnaire, routine laboratory tests, lung function testing, electrocardiogram, and a chest X-ray before the study. There was no evidence of heart or pulmonary disease in any of the subjects. On the day of the study, the volunteers were instructed to avoid alcohol and caffeinated beverages for the 12 preceding hours and to refrain from heavy physical activity since the day before. The local Ethical Committee of biomedical research in Chuvash State University named I. N. Ulyanov approved this study. Informed consent was obtained from all individual participants included in the study.

### 2.2. State anxiety and heart rate variability

All subjects were administered psychophysiological evaluation composed of baseline (rest) and stressor (just before the academic exam) phases. State anxiety (SA) was assessed using State-Trait Anxiety Inventory anxiety scales (Hanin 1976).

Electrocardiogram (ECG) was recorded continuously throughout each experiment via a three-lead ECG at a 1000 Hz sampling rate. Time series of the interbeat intervals RR were extracted automatically from the ECG signal and analyzed using the Heart Rate Variability Analysis Software (HRVAS) (Ramshur 2010).

Time-domain and frequency-domain measures were derived from RR series, including meanRR; the standard deviation of the normal-to-normal (NN) interval series, SDNN; the spectral power of low-frequency (LF: 0.04–0.15 Hz) and high-frequency (HF: 0.15–0.4 Hz) band power and LF/HF ratio. The LF/HF power ratio, although the subject of discussion as to whether it may be used as an index of the sympatho-vagal interaction (Billman 2013), has been widely used in the study of stress (Castaldo et al. 2015; Hamilton and Alloy 2016).

### 2.3 Stress due to university examination

A study of stress reactivity (defined as the difference between exam and rest values) was conducted on a sample of students randomly selected from the original 1,156 subjects. Measures of HRV were natural-log transformed (ln) to meet assumptions of parametric statistics. The sample size was calculated using MedCalc Statistical Software version 17.6 (MedCalc Software bvba, Ostend, Belgium; http://www.medcalc.org; 2017). An admission was made that the study should detect a minimal effect size at a significance level of 0.05 and 80% power (Machin et al. 2011). The sample size was calculated on expected differences (δ) and standard deviations of difference (SD) of HR (δ = 3 beats/min, SD =11.77 beats/min), lnSDNN (δ = 0.09 s, SD = 0.39 s), lnLF (δ = 0.25 ms2, SD = 0.95 ms2), lnHF (δ =0.25 ms2, SD = 1.02 ms2) and lnLF/HF (δ = 0.2, SD = 0.84), on the basis of previous studies from our group (Dimitriev et al. 2008). The minimum sample sizes were estimated to be 123, 148, 116, 133 and 144 for lnHR, lnSDNN, lnLF, lnHF and lnLF/HF, respectively. Therefore, the acceptable sample size was determined to be N 148 for the paired t.

For each variable, the students were divided into three groups, according to the values of the first measurement. The first group of subjects was selected using a baseline measurement of less than the population 25th percentile; the second group represents the middle portion of the distribution, where scores ranged from the 25th percentile to the 75th percentile; and the third group consisted of subjects with baseline measurements greater than the population 75th percentile. Fifteen groups were formed on the basis of HR and HRV, and three groups were formed on the basis of state anxiety. For each variable, students were divided into three groups according to the values of the first measurement: three groups on the basis of HR, three groups on the basis of lnSDNN, three groups on the basis of lnHF, etc. In this way, the same subject could be in different groups depending on the baseline levels of HR and HRV. We analyzed the reaction to stress separately for each group.

### 2.4 Regression to the mean

Ordinary linear-regression analysis was used to identify the RTM. The model was stated as (Follow-up _ Baseline) = constant + *b* × baseline. The RTM effect manifests as a negative correlation between baseline values and stress-induced changes in HR and HRV.

Several studies have shown a strong negative correlation between HR and various indices of HRV (time domain and frequency domain) (Billman et al. 2015) that could affect the interpretation of changes in HRV. To correct the effect of heart rate on HRV indices, we used formulae from our previous study (Dimitriev et al. 2016). Corrected lnLF/HF ratio was calculated from corrected lnLF and corrected lnHF. Then linear-regression analysis was performed again to evaluate the effect of baseline HR and HRV on stress reactivity.

The most widely used model of RTM presupposes that the data are normally distributed. Normality of data distribution was tested with the Shapiro–Wilk W test. A test for normality indicated that the HRV variables (SDNN, LF, HF, LF/HF) required log-transformation to meet the criteria for inclusion in the analysis of RTM.

Estimation of RTM assumes that pretest population means *μ* and variances (σ^2^) are known. Population means and variances were calculated from the general baseline data.

Mee and Chua (1991) proposed a modified paired *t*-test that provides a computationally simple approach to separate RTM and stress effects. This approach involves regression of follow-up measurements *Y*_2_ to (baseline level *Y*_1_ – *μ*):

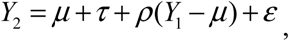

where *τ* is the intervention effect, *ε* is the normally distributed random error, and *ρ* = Corr(*Y*_1_; *Y*_2_). The null hypothesis of no effect is stated as

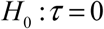

This hypothesis was tested by linear regression analysis of *Y*_2_ – *μ* on *Y*_1_ – *μ*. In this model, the *t*-statistic for testing *τ* = 0 is the *t*-statistic for testing the hypothesis that the intercept equals 0.

### 2.5 Statistical analysis

Repeated-measures analysis of variance (ANOVA) with Bonferroni multiple-comparison test was used to examine the effects of the stress on the HR and HRV measures.

Mean values of change between baseline and follow-up were tested for group differences using univariate analysis of covariance (ANCOVA) with “condition” (< 25th percentile, 25th–75th percentiles, > 75th percentile) as between-subject factors and values of the rest period as covariates.

Means of within-subject distributions were tested against the rest values using a *t*-test for a single sample. All data are presented as mean ± SE. Significance was accepted at *p* < 0.05.

## 3. Results

### 3.1 Stress due to academic examination

The population means of HR, lnSDNN, lnLF, lnHF, and lnLF/HF were 73.11 ± 0.13 bpm, 3.99 ± 0.012 ms, 6.70 ± 0.03 ms^2^, 6.81 ± 0.03ms^2^, and –0.11 ± 0.02, respectively. Table 1 presents the effect of academic stress on HRV measures in three groups.

**Table 1.** Comparison of HRV measures and SA between rest and exam stages

| *Variable* | *All subjects* |  | *1st group*<br>*(< 25th percentile)* |  | *2nd group*<br>*(25th–75th percentile)* |  | *3rd group*<br>*(> 75th percentile)* |  |
| --- | --- | --- | --- | --- | --- | --- | --- | --- |
|  | **Rest** | **Exam** | **Rest** | **Exam** | **Rest** | **Exam** | **Rest** | **Exam** |
| **HR [bpm]** | 73.15 ± 0.89 | 83.49± 0.98# | 60.52 ± 0.60 | 77.52 ± 2.08# | 72.33 ± 0.49 | 81.59 ± 1.11# | 86.96 ± 1.19 | 92.77 ± 1.57* |
| **lnSDNN [ms]** | 3.93 ± 0.03 | 3.78 ± 0.03# | 3.49 ± 0.03 | 3.60 ± 0.05 | 4.03 ± 0.02 | 3.77 ± 0.04# | 4.52 ± 0.04 | 4.13 ± 0.07# |
| **lnLF [ms<sup>2</sup>]** | 6.37 ± 0.07 | 6.21 ± 0.07 | 5.54 ± 0.07 | 5.93 ± 0.12# | 6.69 ± 0.041 | 6.26 ± 0.09# | 7.73 ± 0.06 | 6.94 ± 0.16# |
| **lnHF [ms<sup>2</sup>]** | 6.68 ± 0.09 | 6.03 ± 0.10# | 5.38 ± 0.07 | 5.41 ± 0.18 | 6.81 ± 0.05 | 6.07 ± 0.1# | 8.15 ± 0.08 | 6.78 ± 0.26# |
| **lnLF/HF** | -0.31 ± 0.06 | 0.19 ± 0.06# | -1.17 ± 0.05 | -0.23 ± 0.10# | -0.09 ± 0.04 | 0.36 ± 0.08# | 0.79 ± 0.07 | 0.55 ± 0.14 |
| **SA** | 25.28 ± 0.82 | 41.29 ± 0.83# | 13.48 ± 0.75 | 36.76 ± 1.66# | 24.30 ± 0.49 | 40.86 ± 1.2# | 39.39 ± 0.75 | 46.33 ± 1.66# |

Values are means ± SE.
Exam vs. rest: \* $p < 0.05$ ; # $p < 0.01$

The HF component of HRV was significantly reduced in the exam session compared with the rest session (*p* < 0.001), and the HR ratio was significantly higher in the exam session compared with the rest session (*p* < 0.001)(see Table 1). There was no difference in the LF component of HRV between the exam session and the rest session. lnSDNN reduced significantly from baseline during the exam session (*p*< 0.001).

Academic stress resulted in a significant increase in HR in all three groups. The lnSDNN decreased significantly before the examination in the second and third groups, but not in the group with a low baseline level (*p* > 0.05). The passage from the rest session to the exam evoked a significant increase of lnLF in the first group (*p* < 0.01), whereas lnLF in the other two groups decreased significantly (*F* = 23.46, *p* < 0.01).

Statistical analysis of lnHF revealed a significant decrease in this parameter in the second and third groups and insignificant changes of LnHF were observed in the first group (*F* = 15.39, *p* < 0.01). Analysis of SA changes during the exam session yielded an overall increase in SA. It is clear from Table 1 that the first group has a different pattern of HRV changes.

Results of the population HRV analysis indicate that women had on average a higher heart rate than men (73.94 ± 0.36 vs. 70.59 ± 0.63 bmp, p < 0.01) and lower lnSDNN (3.97 ± 0.01 vs 4.06 ± 0.02 ms, p < 0.01). However, lnSDNN differences disappear after heart rate correction of HRV. Compared to men, women’s HR corrected lnSDNN was 3.11 ± 0.34 versus 3.12 ± 0.34 ms (p > 0.05). LF and HF did not differ significantly by gender (p > 0.05). HRV reactivity to stress was not significantly different between men and women, and stress-related reduction of lnSDNN was similar in both gender groups (men: -0.22 ± 0.11; women -0,15 ± 0.04 ms; p > 0.05). Comparing stress-related decrease in lnHF between women (−0.66 ± 0.11 ms^2^) and men (−0.63 ± 0.30 ms^2^), we found no evidence of a gender difference in stress reactivity (p > 0.05). A similar result was obtained for stress-related reduction of lnLF (men: -0.23 ± 0.2; women: -0.15 ± 0.1 ms^2^, p > 0.05).

### 3.2 Regression to the mean

To reveal potentially significant associations between baseline values and stress effect, we have examined scatter plots of change (exam minus rest measurements) against rest measurement (Barnett et al. 2005) (Fig. 1).

**Figure 1.**
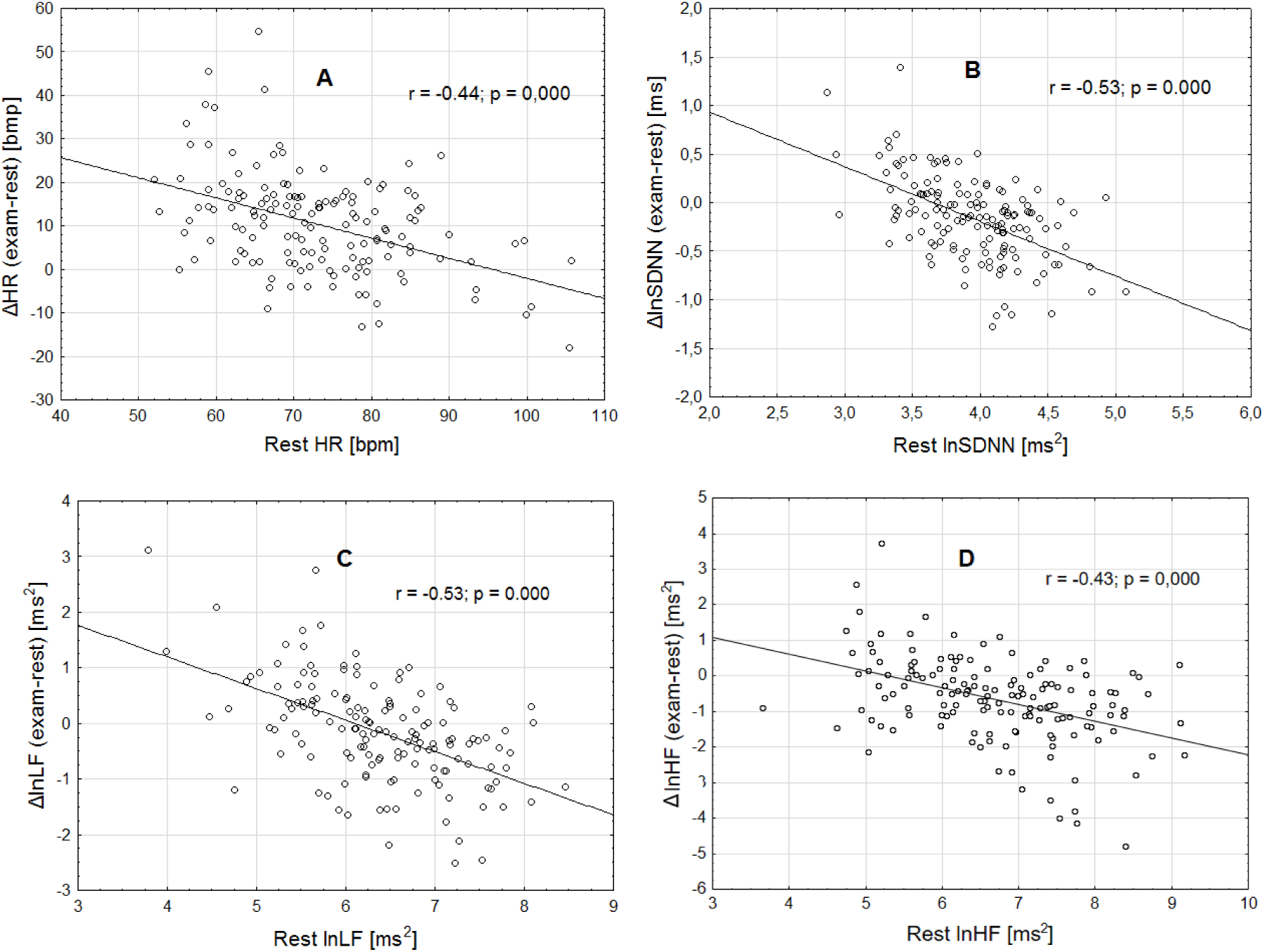
Change (exam – rest) is highly correlated to the baseline level. Scatter plots of baseline HR (A), lnSDNN (B), lnLF (C), and lnHF (D) vs. change. Linear-regression analysis produced significant negative r values for each metric.

Figure 1 shows the changes in HRV measures plotted against baseline levels. For participants, the amount of HRV reduction increased significantly as baseline lnSDNN, lnLF, and lnHF increased (*p* < 0.001 for linear trend for all measures). RTM is evident on the plot, as subjects from the first group have tended to increase HRV, and subjects with very high baseline HRV have tended to decrease lnSDNN, lnLF, and lnHF. The results for baseline and change ln LF/HF display a similar regression tendency (r=-0.56, b=-0.58, p < 0.001).

The changes in HRV, after correction for heart rate, correlated significantly with baseline level (Table 2).

**Table 2.** Associations between baseline and changes in heart rate corrected HRV.

| Variable | Adjusted $R^2$ | $\beta$ for baseline | t | P |
| --- | --- | --- | --- | --- |
| ln SDNN | 0.37 | -0.64 | 9.27 | <0.001 |
| ln LF | 0.33 | -0.61 | 8.62 | <0.001 |
| ln HF | 0.22 | -0.54 | 6.51 | <0.001 |
| ln LF/HF | 0.29 | -0.61 | 7.83 | <0.001 |

To test whether the HRV changes are due to RTM, we applied Mee and Chua’s test. Mee–Chua tests for the HR, lnSDNN, and lnHF confirmed an exam effect (*t* = 12.11, *p* < 0.001, *t* = 6.40, *p* < 0.001, *t* = 7.88, *p* < 0.001, respectively). We find a significant difference in lnLF between the exam and rest session after adjusting for RTM (Mee–Chua *t* = 6.4, *p* < 0.001).

For HR and HRV, the rest level was used as a covariate in an ANCOVA model to estimate the effect of RTM. Table 3 presents the descriptive statistics of change between sessions and the results of the ANCOVA.

**Table 3.** HRV changes between rest and exam (exam – rest) using the ANOVA and ANCOVA

| Variable | Group | ANOVA |  | ANCOVA |  |
| --- | --- | --- | --- | --- | --- |
| | | Mean changes (±SE) | $p^*$ | Mean changes (±SE) | $p^*$ |
| HR [bpm] | 1st group (< 25th percentile) | 17.00 ± 2.05 | < 0.001 | 8.82 ± 2.71 | > 0.05 |
| | 2nd group (25th–75th percentile) | 9.26 ± 1.19 | | 8.66 ± 1.24 | |
| | 3rd group (> 75th percentile) | 5.81 ± 1.63 | | 14.60 ± 2.83 | |
| lnSDNN [ms] | 1st group (< 25th percentile) | 0.11 ± 0.06 | < 0.001 | -0.27 ± 0.10 | > 0.05 |
| | 2nd group (25th–75th percentile) | -0.26 ± 0.04 | | -0.25 ± 0.04 | |
| | 3rd group (> 75th percentile) | -0.39 ± 0.07 | | -0.02 ± 0.11 | |
| lnLF [ms<sup>2</sup>] | 1st group (< 25th percentile) | 0.39 ± 0.12 | < 0.001 | -0.16 ± 0.20 | > 0.05 |
| | 2nd group (25th–75th percentile) | -0.43 ± 0.09 | | -0.41 ± 0.09 | |
| | 3rd group (> 75th percentile) | -0.79 ± 0.15 | | -0.26 ± 0.24 | |
| lnHF [ms<sup>2</sup>] | 1st group (< 25th percentile) | 0.027 ± 0.17 | < 0.001 | -0.39 ± 0.33 | > 0.05 |
| | 2nd group (25th–75th percentile) | -0.74 ± 0.11 | | -0.73 ± 0.13 | |
| | 3rd group (> 75th percentile) | -1.37 ± 0.24 | | -0.97 ± 0.34 | |
| **lnLF/HF** | 1st group (< 25th percentile) | $0.94 \pm 0.10$ | < 0.001 | $0.38 \pm 0.17$ | > 0.05 |
| | 2nd group (25th–75th percentile) | $0.45 \pm 0.09$ | | $0.59 \pm 0.09$ | |
| | 3rd group (> 75th percentile) | $-0.24 \pm 0.14$ | | $0.48 \pm 0.23$ | |

Values are means ± SE.
\*effect of group

When ANCOVA was performed to evaluate the effects of group on changes in HR and HRV, no significant effects were found regarding the changes of HR, lnSDNN, lnLF, and lnHF scores from rest to exam (HR: *F* = 2.03, *p* = 0.13; lnSDNN: *F* = 2.26, *p* = 0.11; lnLF: *F* = 1.14, *p* = 0.32; lnHF: *F* = 0.49, *p* = 0.61, lnLF/HF: *F* = 0.98, *p* = 0.37) after adjusting for the baseline level (Table 3).

The intraindividual distributions of lnHF (Fig. 2) are given below to illustrate the nature of RTM (data from our previous study (Dimitriev DA et al. 2007)). These subjects took part in a study of the relationships between phases of the menstrual cycle and HRV. RR intervals were recorded daily throughout the one month (excluding weekends). The same subjects were examined for acute stress reaction.

**Figure 2.**
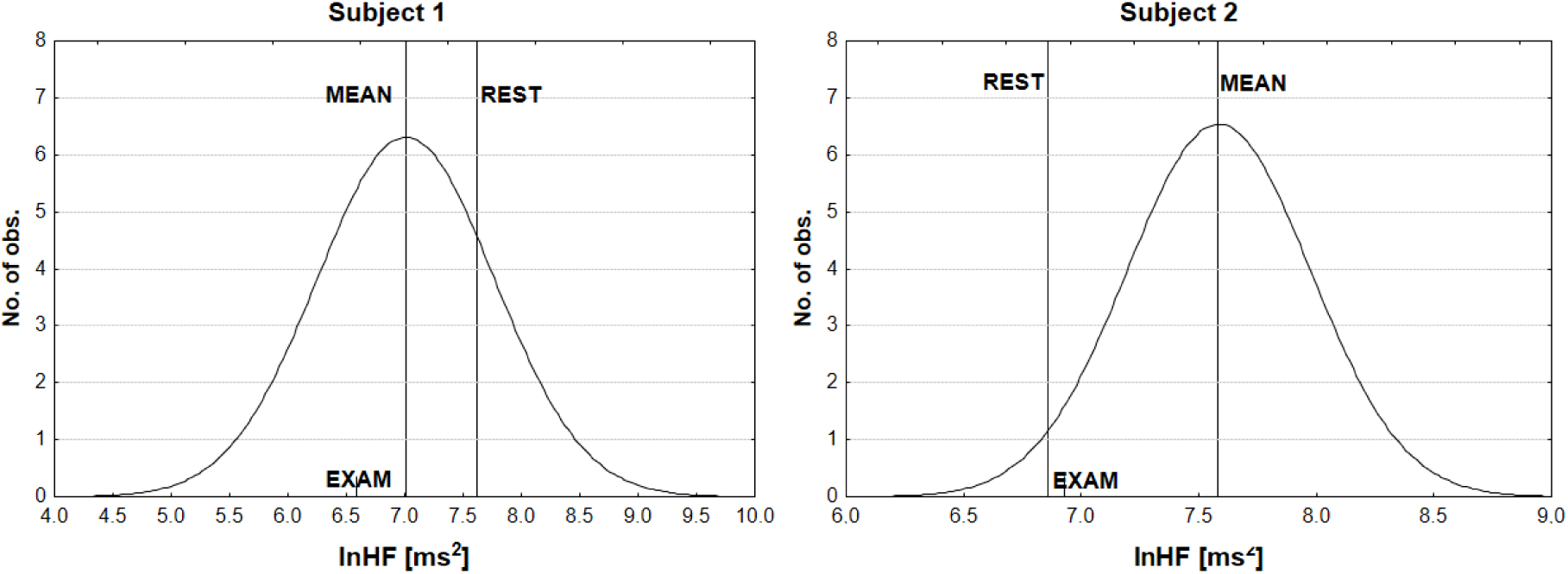
Within-subject distributions and RTM. A mean was calculated from daily lnHF values.

Subject 1‘s lnHF at rest (7.62 ms^2^) was significantly higher than the true mean (7.01 ms^2^; *t* = 7.18; *p* < 0.001). The difference between rest and exam was 1.03 ms^2^ and between mean and exam 0.42 ms^2^. This suggests the stress reaction is overestimated by about 0.61 ms^2^. For the second subject, lnHF at rest (6.86 ms^2^) was significantly lower than the true mean (7.58 ms^2^, *t* = 10.51, *p* < 0.001) and was approximately equal to the lnHF before the exam (6.93 ms^2^). We might think that the subject’s lnHF at exam had slightly increased when in fact the rest measurement was just unusually low, and the subject’s true response was negative (lnHFexam – lnHFmean = –0.65 ms^2^; *t* = 9.49 *p* < 0.01). SA was increased in both subjects (Subject 1: from 28 to 42, Subject 2: from 30 to 38).

## 4. Discussion

A common study design in physiological research is to give subjects some intervention and then observe the response. For example, HRV may be measured twice: once at rest and once during stress. Despite its evident simplicity, the analysis of this form of study may produce biased results due to RTM, although this is not widely appreciated in the literature and has not been quantified. Stress studies in humans reported that stressful life events are associated with a more prominent depression of RSA in individuals with higher baseline HRV, but to our knowledge, none of these analyses included correction for the RTM effect.

A linear regression analysis was used to assess the relation of stress response to baseline HRV value. The results of this analysis show the difference between rest and exam was influenced by baseline level of HRV: subjects with unusually high baseline levels have more prominent stress response than subjects with very low initial values of HRV variables. These findings indicate the strong RTM effect.

Mee–Chua tests for HRV variables confirmed the marked decrease in SDNN and HF. The crude difference in LF between rest and exam almost reached the significance level (*p* = 0.05) and *p*-value drawn from the Mee–Chua test was below 0.025. The authors attribute the observed difference to the RTM effect, leading to LF increasing in the first group, which includes subjects with low baseline levels of LF.

Two analytical strategies have emerged to examine changes between baseline and follow-up measurements of HRV. The first is one-way repeated measures analysis of variance (repeated-measures ANOVA) and the second adjusted for baseline HRV data (ANCOVA). ANCOVA enables removal of the RTM effect and permits estimation of adjusted group means. In this study of HRV reactivity, these two approaches lead to different conclusions: the results of the ANOVA suggest a greater decline of HRV in the third group (very high baseline level) and an increase of HRV variables in the first group (very low baseline level). In contrast, those obtained using ANCOVA with a correction to baseline level suggest the similarity of reactions in all three groups. A source of difference between these two approaches is that ANOVA results are biased by RTM.

RTM is omnipresent in human studies (Gavish et al. 2011; Guenancia et al. 2013; McCambridge et al. 2014; Margaritelis et al. 2016; Forstmeier et al. 2016; Perry and Karpova 2017), and the RTM effect can lead the researcher astray. It most commonly occurs in studies where subjects are selected because they expressed very high or very low baseline values. Thus, participants are eligible to be in the study group only if they have unusually extreme values of RSA or some other variable thought to be related to the stress response. By RTM, subjects with an extremely high level of baseline HRV will have a more prominent stress response, regardless of the severity of the state anxiety. In this case, an individual autonomic flexibility is simply owing to a shift in RSA on the second occasion to the individual mean.

Mee and Chua (1991) presented a modified paired *t*-test that ensures a computationally simple approach to hypothesis testing that accounts for the effect of RTM. The main drawback of this method is that it requires knowledge of the true mean *μ* and variance *σ*^2^ in the target population. In our research population, mean and variance have been estimated from HRV measurements of 1,153 students, and our number of subjects is comparable to those used in other cohort studies of HRV (Kawachi et al. 1995; Tsuji et al. 1996; Sinnreich et al. 1998; Antelmi et al. 2004). However, such data may not always be available, and it is not common to have population data as a basis for stress research protocols. To overcome this problem, Ostermann et al. (2008) developed a straightforward method based on Mee and Chua’s modified *t*-test for the situation, where the population *μ* is unknown.

RTM is caused by random fluctuations in physiological parameters. Random variation is the substantial feature in HR behavior (Laitio 2007; Castiglioni et al. 2009). The HRV can be affected by various factors, such as physical activity, caffeine beverages, menstrual cycle, mood, alertness, mental activity, respiratory rate, and depth. It is easy to control for recreational activity, natural cycles, caffeine consumption, and respiration, but we cannot completely rule out that the person’s emotional and mental state may bias the results of HRV measurements.

The RTM effect can be corrected by adequate study design and/or appropriate adjustment. It has been recognized that multiple baseline measurements could significantly reduce the RTM effect (Barnett et al. 2005). Because of the study design, we could not implement an RTM correction through multiple baseline measurements, which could be regarded as a limitation of our study. We found it to be evident from a wide variety of research papers that the discussion of RTM affects many fields of physiology. Unfortunately, methods to adjust for RTM are rarely used in the analysis of physiological data. We, therefore, believe that the lack of adjustment to RTM in most of the studies probably leads to a significant bias in the results.

## 5. Conclusions

Studies of autonomic flexibility demonstrated that high baseline HRV was associated with a decline in HRV and subjects with low baseline HRV had an inverse reaction to stress, suggesting that baseline HRV is a marker of stress adaptive capacity. The results of our study strongly support an alternative view: RTM is the major source of variability of stress-related changes in HRV. The effect of baseline HRV level disappears after adjustment for RTM.

## Acknowledgments

The study is externally funded by the Ministry of education and science of the Russian Federation (project 19.9737.2017/BCh, http://www.goszadanie.ru). The funder had no role in study design, data collection and analysis, decision to publish, or preparation of the manuscript.

